# The effects of physical and temporal certainty on locomotion with discrete underfoot perturbations

**DOI:** 10.1101/2022.05.04.490667

**Authors:** Nicholas Kreter, Carter Lybbert, Keith E. Gordon, Peter C. Fino

## Abstract

**Background:** Ambulation over complex terrain requires active control of foot placement to maintain a normal kinematic relationship between the center of mass and base of support. Recent investigations have suggested that foot placement location may be selected to anticipate shifts to the underfoot center of pressure. However, it is unclear whether temporal affordance and physical certainty contribute to the selection of a perturbation-specific anticipatory strategy. This study investigates anticipatory and reactive locomotor strategies for repeated underfoot perturbations with varying levels of temporal certainty, temporal affordance, and physical certainty.

**Methods:** Thirteen healthy adults walked with random underfoot perturbations from a mechanized shoe. Temporal certainty was challenged by presenting the perturbations with or without warning. Temporal affordance was challenged by adjusting the timing of a warning tone before the perturbation. Physical certainty was challenged with conditions that included only eversion perturbations, only inversion perturbations, or both eversion and inversion perturbations. Linear-mixed effects models assessed the effect of each condition on the percent change of margin of stability and step width, respectively.

**Results:** For temporally uncertain perturbations and perturbations with one stride or less of affordance, we observed few changes to step width or margin of stability. As affordance increased to two strides, participants adopted a wider step width in anticipation of the perturbation (*p = 0.001*). Physical certainty had little effect on gait for the step of the perturbation, but participants recovered normal gait sooner when the physical nature of the perturbation was predictable (*p < 0.001*).

**Discussion:** Despite having information about the timing and magnitude of upcoming perturbations, individuals do not develop perturbation specific feedforward strategies but instead rely on feedback control to recover normal gait after a perturbation. However, physical certainty appears to improve the efficiency of the feedback controller and allows individuals to recover normal gait sooner.

## 1. Introduction

Bipedal gait requires active neuromotor control on a step-to-step basis to avoid falls (Kuo, Bauby and Kuo, 2000). With each step, the center of mass (CoM) accelerates away from the current stance foot, and in order not to fall, one must redirect the CoM with a new step. It is generally accepted that foot placement is controlled relative to the CoM kinematic state (e.g. its position and velocity) and that this is the dominant mechanism for achieving stable gait (i.e. gait that successfully avoids falls) (Bruijn and Van Dieën, 2018). For example, constraining the CoM during gait creates a reduction in step width (SW), and constraining mediolateral (ML) foot placement creates a similar reduction in ML CoM motion (Arvin *et al*., 2016; Arvin, van Dieën and Bruijn, 2016; Mahaki, Bruijn and Van Dieën, 2019). Furthermore, perturbations to the CoM kinematic state during stance create a predictable shift in the subsequent foot placement (Hof and Duysens, 2013; Wang and Srinivasan, 2014). While the link between the kinematic state of the CoM and foot placement is well known, the relationship between foot placement and disruptions to the center of pressure (CoP) remains less clear.

Complex environments with underfoot perturbations that disrupt the CoP are common to daily living. In such environments, humans use a combination of anticipatory and reactive strategies to maintain balance. For instance, foot placement and joint torque strategies can be used in both feedforward (anticipatory) and feedback (reactive) manners when walking over complex terrain (Bruijn and Van Dieën, 2018; Reimann, Fettrow and Jeka, 2018). Precisely anticipating a gait disturbance requires 1) certainty about when the perturbation will occur, 2) an internal representation of the physical disturbance (e.g., location and magnitude relative to one’s own body state), and 3) enough time to develop and execute a motor command (Figure 1). When individuals lack certainty about the physical nature or specific timing of an upcoming disturbance, a cautious gait strategy may be adopted, characterized by a shorter step length and wider SW. For example, healthy young adults have been shown to exhibit cautious gait when walking while blindfolded (Hallemans *et al*., 2009; Saucedo and Yang, 2017), and when they are generally aware of upcoming slip or translation perturbations but don’t know specific timing (Lawrence *et al*., 2015; Nestico *et al*., 2021). Yet, it remains unclear how the temporal and physical nature of upcoming perturbations contribute to cautious gait; perturbations with certain temporal and physical features, and enough temporal affordance, may allow individuals to implement a precise anticipatory strategy specific to task demands.

**Figure 1:**
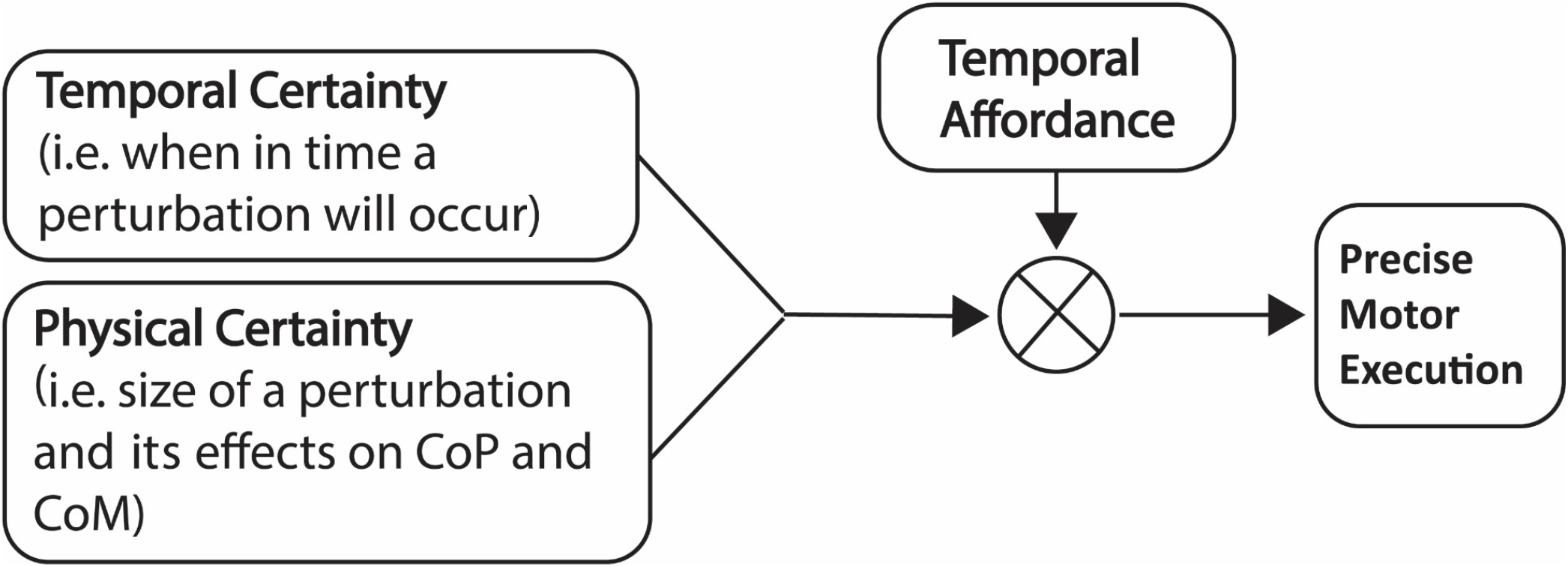
Factors for anticipating gait disturbances. To implement a task-specific anticipatory strategy, one needs temporal and physical certainty to know when in time the disturbance will occur and how it will impact their gait. If the temporal and physical qualities of a perturbation are certain, the remaining factor that could impede a precise response is whether there is the level of temporal affordance (i.e., enough time) between knowing a disturbance will occur and experiencing the disturbance.

We previously reported results in which healthy young adults completed walking tasks with underfoot eversion perturbations similar to stepping on a small rock that were either unexpected (no warning) or expected (warning tone approximately 600 ms before the perturbation) (Kreter, Rogers and Fino, 2021). When perturbations were unexpected, participants exhibited no kinematic differences relative to their normal walking pattern. Yet, when given a warning tone one stride prior to a perturbation, participants exhibited a different strategy in the swing phase prior to foot placement. Specifically, the acceleration profile of the swing foot suggested that they adopted a more medial foot placement relative to their normal gait. In theory, a medial shift to foot placement would neutralize the lateral shift to the CoP from the perturbation and allow participants to retain a normal kinematic relationship between the CoP and CoM. We speculated that participants may have developed an internal model of the physical characteristics of the perturbation through sensory feedback and implemented an anticipatory strategy specific to the task’s physical demands. However, the reliance on acceleration data made it difficult to conclusively characterize the anticipatory strategies that participants adopted.

The purpose of this study was to further probe how healthy adults anticipate well-defined CoP perturbations with differing levels of physical certainty, temporal certainty, and temporal affordance. Temporal certainty was challenged by delivering an underfoot perturbation with or without advanced warning, temporal affordance was challenged by providing warnings of upcoming disturbances with 0.5, 1, and 2 strides notice before the perturbation, and physical certainty was challenged by randomly delivering perturbations that either inverted or everted the foot. We hypothesized that individuals would adopt a perturbation-specific anticipatory stepping strategy, seeking to minimize the disruption to the normal kinematic relationship between the CoM and CoP, if they knew when (temporal certainty) to expect the perturbation and how (physical certainty) it would disrupt their CoP. We also hypothesized that anticipatory stepping strategies would be limited as the temporal affordance decreased. Finally, we expected that reducing an individual’s certainty related to the physical characteristics of a perturbation would lead to a more cautious gait strategy featuring a larger margin of stability (MoS) and wider SW before the perturbation.

## 2. Materials and Methods

### a. Participants

A power analysis (G*Power, Universitat Kiel, Germany) with an effect size of (*f = 0.345*), power threshold of 0.8, and a significance level of *a* = 0.05 revealed a sample size of 13 participants would be appropriate for this study. An effect size (*f = 0.69*) was calculated using data from our previous study that had trials with unexpected (zero affordance) and expected (one stride affordance) perturbations in healthy control participants. Due to the more specific levels of affordance tested here, this project was powered using an effect size that was 50% of the magnitude of the one above (*f = 0.345*).

Thirteen healthy young adults [26.8 (5.6) years, 6 female] provided written informed consent in this IRB approved study. Exclusion criteria included (1) a history of neurological or behavioral pathologies that could explain balance deficits, (2) a history of musculoskeletal injuries within the past year that could impact balance, (3) any reconstructive surgery of the lower extremities, (4) and any current medications that could cause neuromotor deficits.

### b. Protocol

Participants wore a set of custom shoes containing small, mechanized plastic blocks just proximal to the 1^st^ and 5^th^ metatarsophalangeal joints of the left foot (Figure 2). The blocks were controlled by micro-servo motors housed in the sole of the shoe. The block and motor were recessed into the sole such that they wouldn’t interfere with normal gait (Kim *et al*., 2013). When triggered, the recessed block deployed during the swing phase of gait and resulted in a ~6° inversion of eversion of the ankle. A pair of force sensitive resistors on the heel of the shoe communicated with a microcontroller attached to the side of the shoe and registered heel strikes. For trials with perturbations, the control box would trigger the perturbation randomly between every fifth and ninth stride. During most trials participants received a warning that the perturbation was upcoming. The warning came in the form of a loud beep, played through a pair of Bluetooth headphones that the participants wore as they walked (Kreter, Rogers and Fino, 2021). Participants were informed that the timing of the perturbation warning would remain consistent within trials but may change between trials.

**Figure 2:**
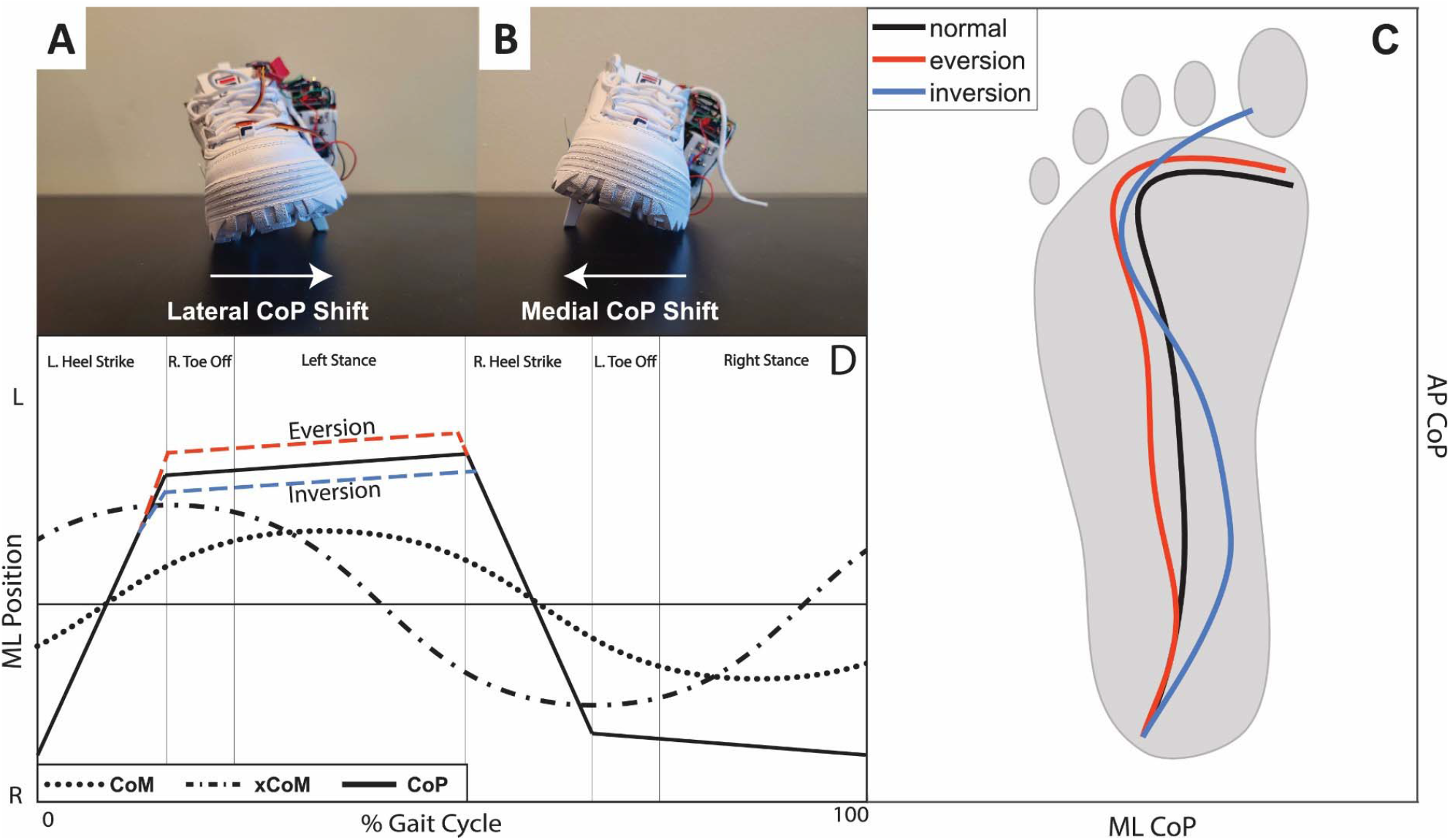
The mechanized shoe. Participants completed walking trials with repeated eversion (A) and inversion (B) perturbations from the motorized flaps housed in the sole of the shoe. Participant CoP data (C) show that the CoP is shifted towards the lateral aspect of the foot for eversion perturbations and towards the medial aspect of the foot for inversion perturbations. (D) Participant data displaying the shift to CoP associated with each perturbation. Underfoot shifts to CoP may disrupt the normal kinematic relationship between the xCoM/CoM and CoP during stance phase of gait. However, if the perturbation is familiar and can be anticipated, stepping behavior may be adapted to maintain a normal relationship.

While wearing the shoes, participants completed eight walking trials, six of which contained perturbations. Participants walked along an instrumented treadmill at a comfortable self-selected speed. Participants spent 10 minutes to acclimate to treadmill walking before any trials were recorded. To determine preferred walking speed, participants started walking at a slow speed that was increased in increments of 0.01 m/s until they reported they were at their preferred speed. The speed was then increased by 0.4 m/s and decreased in increments of 0.01 m/s until the participant again reported that they had reached their preferred walking speed. This process was repeated until two consecutive reports of preferred walking speed were within 0.05 m/s (Jordan, Challis and Newell, 2007).

Participants completed six walking trials with perturbations, each five minutes in length, and a pair of two-minute trials without perturbations before and after the perturbation trials. The first perturbation trial was always an unexpected condition that included perturbations and no warning. The remaining five perturbation trials were split into two sequences that manipulated either temporal affordance or physical certainty. The temporal affordance sequence contained three randomized expected perturbation conditions with warning tones occurring one half of a stride (i.e., one step), one stride (i.e., two steps), and two strides (i.e., four steps) before the perturbation, respectively. The second sequence contained two additional conditions. The first involved expected inversion perturbations with a one stride warning. The second contained random delivery of inversion or eversion perturbations with a one stride warning. Before the first perturbation trial (unexpected condition), participants were informed that the following six trials would involve small underfoot perturbations that felt like stepping on a small rock. After the unexpected perturbation trial, participants were told that for the remaining perturbation trials they would hear warning tone before the perturbation occurred. Each perturbation trial included around 30 perturbations (mean = 31.9, std = 6.7 perturbations).

**Figure 3:**
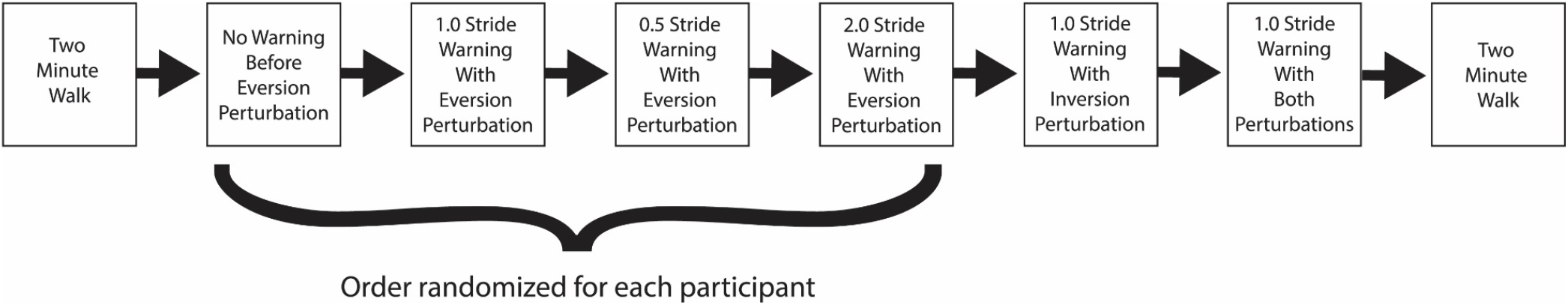
Walking trial progression. Participants will complete a single task walk, followed by a sequence of four trials that constrain response time. The remaining trials will introduce the inversion perturbation then randomly switch between inversion and eversion to test how anticipatory strategy changes with less physical certainty.

### c. Instrumentation

A 3D motion capture system (Bonita V10, VICON, Oxford, UK) surrounding the testing environment captured positional marker data at a rate of 200 Hz. Kinetic data were recorded at a rate of 1000 Hz from two force plates embedded beneath the left and right belts of an instrumented treadmill (BERTEC, Columbus, OH). Participants were outfitted with reflective markers in a custom lower-body marker configuration. The custom marker configuration included markers bilaterally at the anterior superior iliac spine (ASIS), posterior superior iliac spine (PSIS), the lateral epicondyle of the femur, and the lateral malleolus. Four markers were placed on each shoe including at the 1^st^, 2^nd^, and 5^th^ metatarsophalangeal joints, and the calcaneus. Forces, moments, and CoP position were recorded from the force platforms embedded under each belt of the treadmill. Each force platform was oriented so that the positive y-axis is aligned with forward progression and the positive x-axis points to the left.

### d. Outcome Measures

Lateral MoS (Hof, Gazendam and Sinke, 2005) and SW were calculated for each step of each trial as measures of gait stability. MoS is a measure of dynamic stability that assesses the distance between the extrapolated center of mass (xCoM) and the base of support (BoS) (Hof, 2008; Watson *et al*., 2021). The x-position of the CoP was used to approximate the lateral BoS (BoS). The geometric center between the four reflective markers of the pelvis was used to estimate the position of the CoM. The velocity of the CoM (vCoM) was calculated using a central difference algorithm. To account for the lack of forward progression during treadmill walking the velocity of the treadmill belt was added to the vCoM (Fallahtafti *et al*., 2020). For this study, the lateral direction was defined using the global reference frame aligned with the treadmill. However, to clarify interpretation of MoS, the positive direction was opposite for left and right steps. MoS was identified at contralateral toe-off for each step and SW calculated as the difference between contralateral heel strikes in the ML direction. The percent change in MoS and SW were calculated relative to the mean of normal steps within a given trial. Steps were classified as normal if they were not within the four steps preceding or following a perturbation.

### e. Data Processing and Statistical Analysis

Motion capture data was processed in Vicon Nexus (ver. 2.12). Positional marker data were checked for marker switching and a rigid body fill was used to plug missing gaps. Gait events, outcome measures, and statistical tests were computed with a custom code in MATLAB (ver. R2020b, The Mathworks Inc., Natick MA, USA). Kinetic and kinematic data were filtered using a 4^th^ order phaseless low-pass Butterworth filter. Heel strike and toe off events were determined with an algorithm using the maximal displacement of the heel and toe markers from the sacrum (Zeni, Richards and Higginson, 2008). Perturbation steps were identified by the change in foot roll angle during stance. For statistical analysis, two anticipatory steps, the perturbation step, and three recovery steps were isolated and compared against the inter-trial unperturbed steps. To account for participants acclimating to the association between the warning tone and perturbation in each trial, the first three perturbations in each trial were excluded from data analysis. The number of steps to return to normal gait following the perturbation was also determined using the 95% confidence interval of the percent difference between perturbation steps and normal steps.

To test the effects of temporal certainty, temporal affordance, and physical certainty on balance measures within subjects, linear mixed-effects models were fit with fixed effects of condition and random intercepts for subjects. A test of fixed effects assessed significant differences between conditions. Pairwise contrasts were performed for conditions that reported a significant test of fixed effects. Supplementary correlation analyses were performed to assess the association between MoS during the perturbation step and SW at the first recovery step. All statistical tests were assessed at an alpha level of 0.05.

## 3. Results

### a. Effect of Temporal Affordance and Temporal Certainty on MoS and SW

Changes to temporal affordance and temporal certainty had no statistically significant effects on MoS for any steps surrounding the perturbation with the exception of the second step after the perturbation (Figure 4). For the second step after the perturbation, there was a significant effect of condition on MoS (*p = 0.047*). Pairwise contrasts confirmed the significant effect of affordance, revealing significant difference between the condition with two strides of warning and the condition with one stride (*p = 0.006*). There were no significant effects of certainty on MoS for any step.

**Figure 4:**
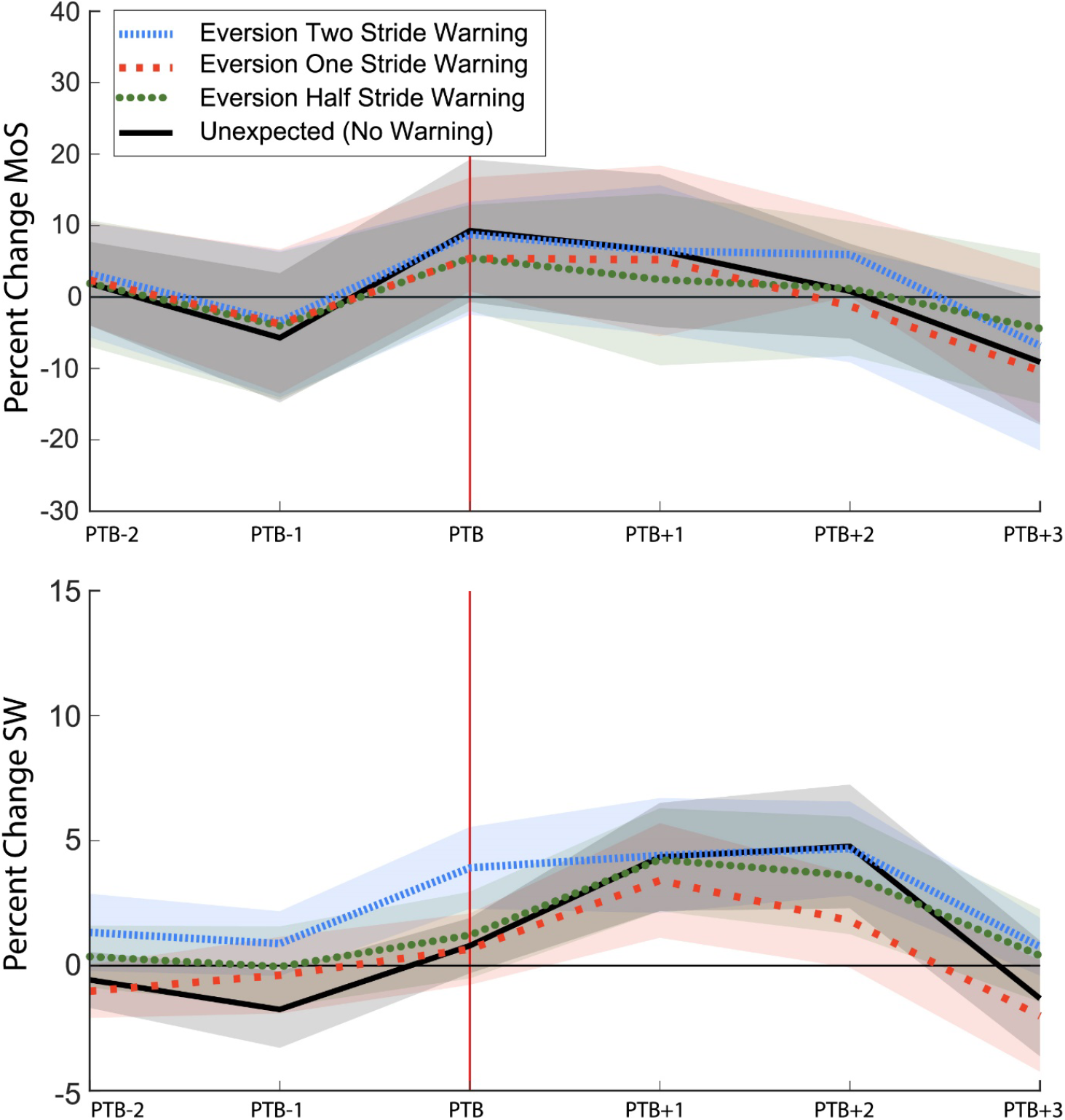
Percent change to MoS (top) and SW (bottom) for conditions with two strides (blue, short-dash), one stride (red, long-dash), a half stride (green, dotted), and unexpected (black, solid) conditions. Solid lines represent the mean and shaded regions represent the 95% confidence intervals for each condition. Gait was considered normal when the 95% confidence interval crossed zero for a given step.

**Table 1:** Percent change to MoS and SW relative to unperturbed gait for trials testing the effects of temporal certainty and temporal affordance. Means and 95% confidence intervals are presented for two anticipatory steps, the perturbation step (PTB) and three recovery steps of each condition

| Condition | PTB-2 | PTB-1 | PTB | PTB+1 | PTB+2 | PTB+3 |
| --- | --- | --- | --- | --- | --- | --- |
| **MoS ‡** |  |  |  |  |  |  |
| **Eversion Two** | 3.2 (-4.0 – 10.4) | -3.4 (-13.5 - 6.7) | 8.8 (0.8 – 16.7) | 6.5 (-5.5 – 18.4) | **5.9 (0.1 – 11.8)<sup>‡</sup>** | -6.8 (-17.6 – 4.0) |
| **Eversion One** | 2.4 (-5.6 – 10.4) | -3.8 (-14.0 – 6.5) | 5.4 (-2.5 – 13.3) | 5.2 (-5.2 – 15.6) | **-1.3 (-9.1 – 6.6)<sup>†</sup>** | -10.3 (-21.5 – 0.7) |
| **Eversion Half** | 1.9 (-6.9 – 10.7) | -4.1 (-14.4 – 6.3) | 5.5 (-1.9 – 12.9) | 2.4 (-9.6 – 14.5) | **1.2 (-8.2 – 10.6)<sup>†</sup>** | -4.4 (- 14.9 – 6.1) |
| **Unexpected** | 1.9 (-4.0 – 7.7) | -5.7 (-14.8 – 3.3) | 9.3 (-0.7 – 19.3) | 6.5 (-4.2 – 17.2) | 0.8 (-5.8 – 7.4) | -9.1 (-17.9 – -0.3) |
| **SW** |  |  |  |  |  |  |
| **Eversion Two** | **1.3 (-0.2 – 2.9)<sup>#</sup>** | 0.9 (-0.4 – 2.2) | **3.9 (2.3 – 5.5)<sup>#, ‡</sup>** | 4.4 (2.1 – 6.7) | **4.7 (2.8 – 6.6)<sup>#</sup>** | **0.8 (-0.4 – 1.9)<sup>#</sup>** |
| **Eversion One** | **-1.0 (-2.1 – 0.1)<sup>†</sup>** | -0.4 (-1.9 – 1.1) | **0.7 (-0.8 – 2.1)<sup>†</sup>** | 3.5 (1.1 – 5.7) | **1.8 (-0.1 – 3.7)<sup>†</sup>** | **-2.0 (-4.2 – 0.28)<sup>†</sup>** |
| **Eversion Half** | 0.4 (-0.9 – 1.6) | -0.0 (-1.7 – 1.6) | **1.2 (-0.5 – 3.0)<sup>†</sup>** | 4.2 (2.2 – 6.3) | 3.6 (1.3 – 6.0) | **0.4 (-1.4 – 2.3)<sup>#</sup>** |
| **Unexpected** | -0.6 (-1.7 – 0.5) | **-1.8 (-3.3 – -0.2)<sup>¥</sup>** | **0.8 (-0.3 – 1.9)<sup>¥</sup>** | 4.3 (2.2 – 6.5) | 4.8 (2.3 – 7.2) | -0.3 (-3.6 – 1.0) |
| **† = significantly different from eversion two, # = significantly different from eversion one, ‡ = significantly different from eversion half, ¥ = significantly different from temporally certain condition** |  |  |  |  |  |  |

Affordance had a significant effect on SW at two steps before the perturbation (*p = 0.046*), the perturbation step (*p = 0.005*), the second recovery step (*p = 0.009*) and the third recovery step (*p = 0.015*). Pairwise contrasts confirmed the significant effects at each step. Specifically, two steps before the perturbation participants had an increased step width in the two-stride warning condition relative to the one stride warning condition (*p = 0.009*). During the step with the perturbation, participants had a larger step width for the two-stride warning condition relative to all other levels of affordance (one-stride *p = 0.002*, half-stride *p = 0.008*). At the second recovery step only the one-stride warning condition returned to a normal SW and was significantly less than the two-stride warning condition (*p = 0.004*). By the third recovery step all conditions returned to normal, however, the two-stride and half-stride warning conditions were still significantly elevated relative to one-stride warning condition (two-stride *p = 0.006*, half-stride *p = 0.016*) (Figure 4). Contrasts revealed a significant effect of temporal certainty on SW in the step before the perturbation (*p = 0.024*) and the step of the perturbation (*p = 0.009*).

### b. Effect of Physical Certainty on MoS and SW

No significant effects to MoS were observed during the preparatory or perturbation steps (Figure 5). There was a significant effect of condition on MoS during the second (*p = 0.014*) and third (*p = 0.004*) recovery steps following the perturbation. For the second recovery step there was a significant difference between the mixed perturbation and eversion perturbation conditions (*p = 0.004*). For the third recovery step the mixed perturbation condition was significantly elevated relative to both eversion (*p = 0.002*) and inversion (*p = 0.015*) conditions.

**Figure 5:**
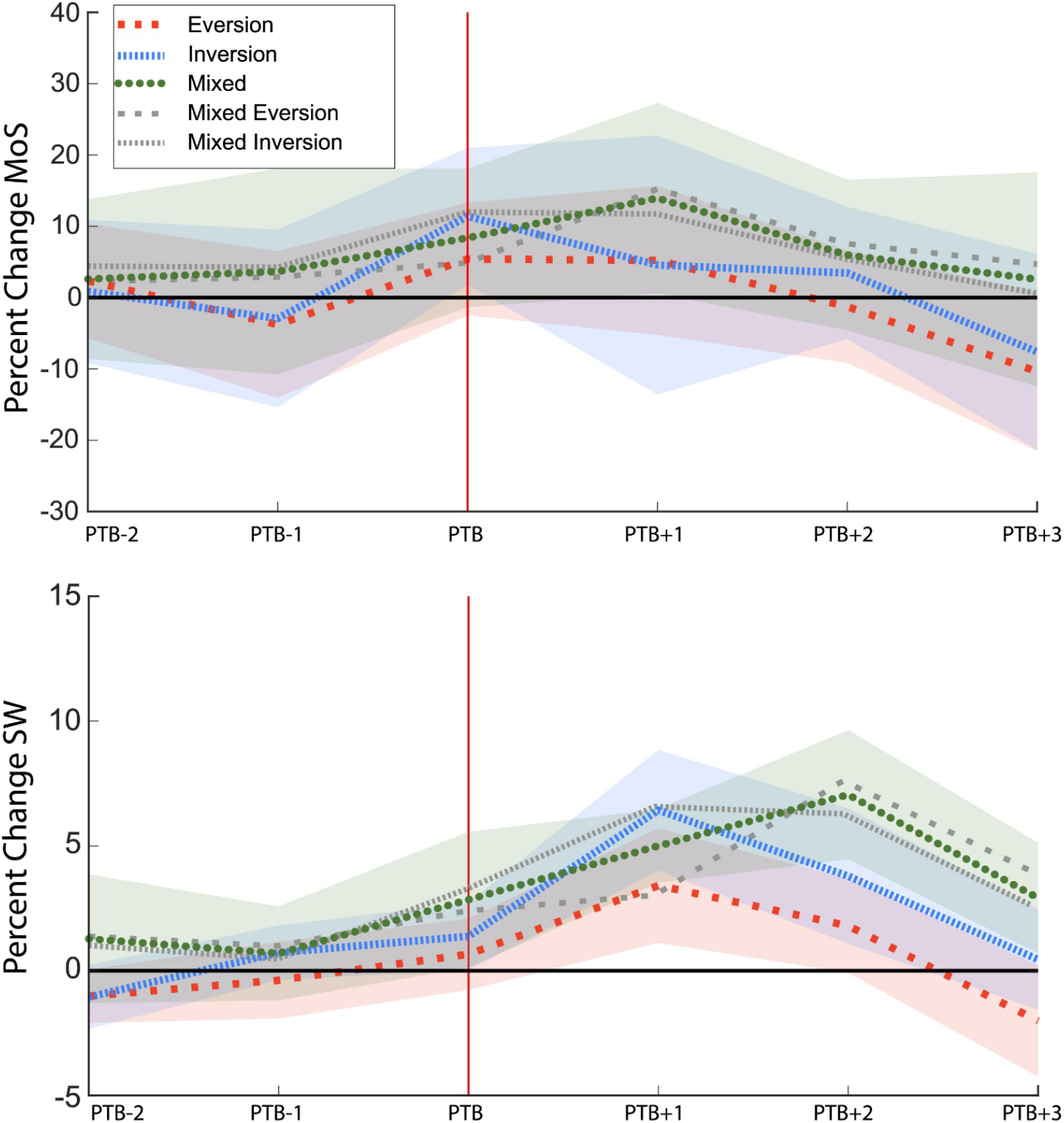
Percent change to MoS (top) and SW (bottom) for eversion only (red, long-dash), inversion only (blue, short-dash), and mixed (green, dotted) perturbation conditions. Grey lines surrounding the mixed perturbation condition represent the means for eversion (long-dash) and inversion (short dash) perturbations within the mixed perturbation condition. Solid lines represent the mean and shaded regions represent the 95% confidence intervals for each condition.

**Table 2:** Percent change to MoS and SW relative to unperturbed gait for trials testing the effects of physical certainty. Means, standard deviations and 95% confidence intervals are presented for the perturbation step (PTB) and three recovery steps of each condition.

|  |  |  |  |  |  |  |
| --- | --- | --- | --- | --- | --- | --- |
| **Condition** | **PTB-2** | **PTB-1** | **PTB** | **PTB+1** | **PTB+2** | **PTB+3** |
| **MoS** |  |  |  |  |  |  |
| **Eversion** | 2.4 (-5.6 – 10.4) | -3.8 (-14.0 – 6.5) | 5.4 (-2.5 – 13.3) | 5.2 (-5.2 – 15.6) | **-1.3 (-9.1 – 6.6)<sup>‡</sup>** | **-10.3 (-21.5 – 0.7)<sup>‡</sup>** |
| **Inversion** | 0.6 (-9.5 – 10.7) | -2.7 (-15.2 – 9.8) | 11.0 (1.2 – 20.7) | 4.3 (-13.8 – 22.3) | 3.2 (-6.2 – 12.6) | **-7.3 (-21.8 – 6.5)<sup>‡</sup>** |
| **Mixed** | 2.5 (-8.7 – 13.7) | 3.2 (-10.9 – 17.3) | 8.0 (-1.8 – 17.7) | 14.7 (1.1 – 28.3) | **5.9 (-4.6 – 16.5)<sup>#</sup>** | **1.8 (-12.9 – 16.5)<sup>#,†</sup>** |
| **SW** |  |  |  |  |  |  |
| **Eversion** | -1.0 (-2.1 – 0.1) | -0.4 (-1.9 – 1.1) | 0.7 (-0.8 – 2.1) | **3.5 (1.1 – 5.7)<sup>†</sup>** | **1.8 (-0.1 – 3.7)<sup>‡</sup>** | **-2.0 (-4.2 – 0.28)<sup>†,‡</sup>** |
| **Inversion** | -1.3 (-2.6 – -0.1) | 0.7 (-0.4 – 1.8) | 1.4 (0.1 – 2.8) | **6.3 (3.9 – 8.7)<sup>#</sup>** | **3.5 (0.7 – 6.3)<sup>‡</sup>** | **0.5 (-1.4 – 2.5)<sup>#</sup>** |
| **Mixed** | 0.9 (-1.4 – 3.3) | 0.5 (-1.2 – 2.1) | 2.4 (0.4 – 4.4) | 4.9 (3.5 – 6.3) | **6.9 (4.4 – 9.4)<sup>#,†</sup>** | **2.7 (0.7 – 4.8)<sup>#</sup>** |
| **# = significantly different from eversion, † = significantly different from inversion, ‡ = significantly different from mixed** |  |  |  |  |  |  |

There was a significant effect of physical certainty on SW during the each of the three recovery steps (recovery step one (*p = 0.044*), recovery step two (*p = 0.009*), and recovery step three (*p < 0.001*)). For the initial recovery step, pairwise contrasts revealed that SW was significantly wider for the inversion condition than the eversion condition (*p = 0.013*), but not the mixed condition (*p = 0.213*). During the second recovery step, participants exhibited a significantly wider SW during the mixed condition than both the eversion (*p = 0.002*) and inversion (*p = 0.039*) conditions. During the third recovery step, SW was still significantly wider during the mixed perturbation condition relative to the eversion condition (*p < 0.001*), but not the inversion condition (*p = 0.058*) (Figure 5).

### c. Relationship Between MoS During Perturbation and SW During Recovery

MoS during the perturbation step was a significant predictor of SW during the recovery step for all conditions (Figure 6). The MoS during eversion with one stride of warning explained the least amount of variance in SW during the first recovery step (*r^2^ = 0.22, p < 0.001*) and the MoS during inversion with one stride of warning exhibited the strongest relationship (*r^2^ = 0.43, p < 0.001*). For trials testing the effect of temporal affordance, MoS during the perturbation step explained 32% of the variance in SW during the first recovery step (*r^2^ = 0.32, p < 0.001*). For trials testing the effect of physical certainty, MoS during the perturbation step explained 32% of the variance in SW during the first recovery step (*r^2^ = 0.32, p < 0.001*).

**Figure 6:**
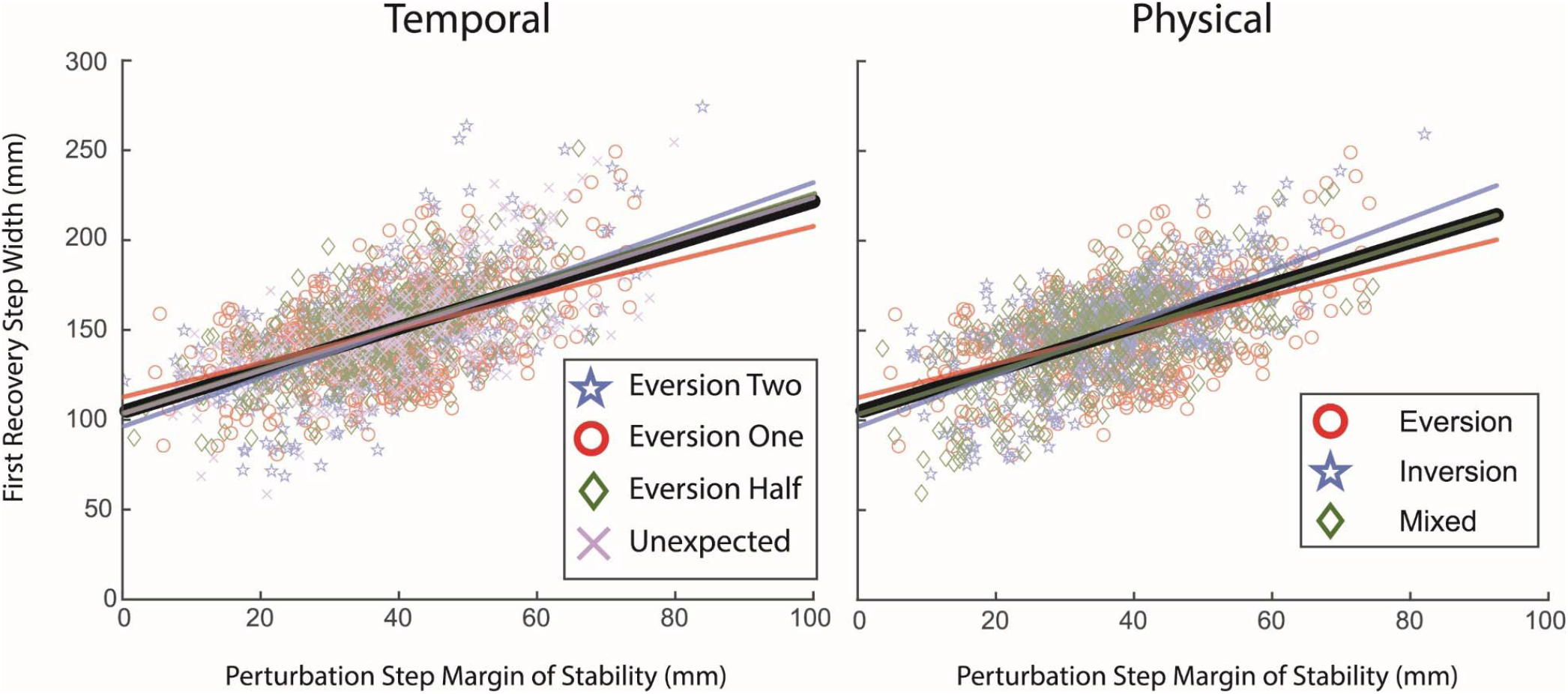
Linear fits between MoS during a perturbation steps and SW of the first recovery step. The left panel includes trials that tested the effect of temporal certainty or affordance and the right panel includes trials that tested the effect of physical certainty. For trials where temporal affordance was manipulated MoS explained 32% of the variance in SW (thick black line, left panel) and for trials where physical certainty was manipulated MoS explained 32% of the variance in SW (thick black line, right panel). Scatter points display each perturbation for each trial across all participants, separated by trial.

## 4. Discussion

This study aimed to further investigate the effects of (1) temporal certainty, (2) temporal affordance, and (3) physical certainty on normal kinematics during gait with expected underfoot perturbations. We predicted that, with greater temporal certainty, participants would select a specific foot placement to counteract the shift to CoP from the underfoot perturbation, but that this strategy would become less feasible, and thus less frequent, as temporal affordance decreased. For this perturbation, we did not find a significant effect of temporal certainty or temporal affordance except when two strides of warning were provided. For steps with greater physical certainty, we expected participants to implement a task-specific foot placement strategy to mitigate changes to underfoot CoP, but adopt a cautious strategy when the perturbation was less physically certain. Specifically, we expected a wider SW for inversion perturbations and a narrower SW for eversion perturbations, but no change to MoS. We did not observe perturbation-specific foot placement strategies with greater physical certainty; however, participants required more steps to recover their normal gait when perturbations had less physical certainty.

### a. Temporal Certainty and Affordance Had Little Impact on Anticipatory or Recovery Strategies

Counter to our hypothesis, we did not observe a reduced SW before temporally certain eversion perturbations, regardless of temporal affordance. The most noteworthy change to stepping behavior occurred during the two-stride warning condition, where participants adopted a wider SW in anticipation of the perturbation. One explanation for the increase with two strides of warning is that the warning may have been delivered too early. Potential stepping locations are typically sampled during the stance phase one stride prior to heel contact, and if visual information for stepping locations are available outside of this time frame, but not during, stepping error increases (Matthis and Fajen, 2014; Matthis, Barton and Fajen, 2017). Planning foot placement one stride in advance is biomechanically advantageous because it allows humans to fine-tune the balance between energetically efficient gait and stability. More explicitly, it allows for the use of foot placement in the preceding step to tailor the location of the BoS while the trailing limb can adjust push-off force and change CoM trajectory (Kuo and Donelan, 2010). Planning steps too far in advance doesn’t necessarily have a biomechanical disadvantage, but it does require an individual to store a larger number of prepared foot placement locations in their working memory. In object manipulation tasks, humans minimize the use of visual working memory until it is necessary to complete the task at hand (Ballard, Hayhoe and Pelz, 1995). In a stepping task such as ours, storing the information for a specific foot placement four steps ahead (i.e. the two-stride warning condition) while also planning the three preceding steps may have been undesirable, leading participants to adopt a less effective anticipatory strategy instead.

The absence of a clear anticipatory strategy may also be due to the participants’ inability to use vision to anticipate the disturbance. Foot placement is primarily guided by visual information during locomotion (Gibson and Crooks, 1938; Patla, 2004), and individuals may track obstacles until they step on them in the real world. Here, participants were forced to create an internal model of the perturbation’s physical qualities by integrating proprioceptive information over repeated exposures. Participants had no reason to direct foot placement through vision as the disturbance was unavoidable and couldn’t be seen. Given the futility of anticipating foot placement through vision, participants may have instead focused on developing an effective recovery strategy reliant on proprioceptive feedback.

### b. Physical Certainty of Upcoming Perturbations Allows for Quicker Recovery of Stability

Counter to our hypotheses, there were no clear anticipatory adjustments to MoS or SW for the steps leading up to the perturbation. However, participants in our study achieved a normal SW sooner when the perturbation had predictable physical features than when it randomly switched between inversion and eversion. Notably, when inversion and eversion perturbations from the mixed-condition were assessed separately, the changes to MoS during the perturbation step and SW during the first recovery step mirrored the changes observed in the physically certain trials (Figure 5). The quicker recovery during trials with predictable perturbations suggests that individuals are able to form a physical representation of the perturbations features and prime a response that allows them to better anticipate how a perturbation will disrupt their gait and recover more effectively. Groups with deficits to sensory integration may struggle to recover normal gait around similar perturbations in everyday life. When exposed to perturbations from a similar shoe mechanism, persons with peripheral neuropathy struggled to modify step length in response to unexpected inversion perturbations (Allet *et al*., 2014), and older adults adopt a hip torque strategy (Kim *et al*., 2013), typically reserved for larger disturbances. Future work should focus on groups with neuromotor deficits and incorporate neurological recordings such as electroencephalography (EEG) to better delineate the formation and execution of motor plans in anticipation of known upcoming disturbances (Nordin, Hairston and Ferris, 2019).

### c. Feedforward/Feedback Controllers

When planning foot placement during locomotion, humans incorporate both feedforward and feedback control strategies to optimize efficiency and stability (Bruijn and Van Dieën, 2018). Selection of feedforward or feedback strategies is dependent on a multitude of variables, including complexity of the walking environment and certainty related to upcoming disturbances. Feedforward locomotor strategies allow humans to anticipate gait disturbances by predicting a future body-state while feedback strategies are used to assess instantaneous body state to recover a normal state after a disturbance. We hypothesized that greater temporal and/or physical certainty related to upcoming perturbations would allow participants to adopt a specific feedforward strategy that incorporated proprioceptive information from repeated exposures to our perturbation to predict the effect of upcoming perturbations. Yet, our results suggest that participants primarily relied on a feedback control strategy to manage frontal plane stability. Specifically, the disruption to MoS during the perturbation step influenced SW during the first recovery step (Figure 6). Even for the condition with two strides of warning where participants did exhibit feedforward control prior to the perturbation, MoS during the perturbation step was still a good predictor of SW in the first recovery step. These results are congruent with studies showing that instantaneous CoM dynamics guide foot placement in the subsequent step (Arvin, van Dieën and Bruijn, 2016; Mahaki, Bruijn and Van Dieën, 2019).

Despite greater temporal affordance failing to demonstrably aid perturbation recovery, greater physical certainty allowed participants to implement a more effective feedback strategy and return to normal gait after just one or two recovery steps. The use of a feedback control strategy may be due to the infinite number of body-states an individual can adopt to accomplish a step with a perturbation. While step-to-step kinematic differences are subtle, any change creates a novel kinematic state to predict a disturbance within – our previous study with a similar perturbation demonstrated different magnitudes of ankle roll for different stepping characteristics (Kreter, Rogers and Fino, 2021). Even though the disturbance in this study has fixed physical qualities, predicting its exact impact on body state with a feedforward model is less certain than waiting to develop a feedback model, especially given the amount of time individuals have between each step (~500 ms). Other recent studies with predictable and unpredictable shifts to the CoM during gait also suggest that individuals use feedback from instantaneous shifts to the kinematic state to recover stability and that certainty related to the timing or direction of a disturbance may just enhance the feedback control strategy (Major, Serba and Gordon, 2020). Additionally, the perturbations used in our study are very small and pose little threat to stability for healthy adults; future work should investigate perturbations of greater magnitude or perturbations with a faster cadence as these factors may force individuals to weigh feedforward and feedback control strategies differently.

### d. Limitations

This study should be interpreted within the context of its limitations. First, only the left foot received perturbations. A unilateral perturbation was selected to simplify the interpretation of the warning. Participants may have adopted gait strategies to support the left limb and mitigate the impact of perturbations. Such strategies may have included anticipatory strategies that were not measured in this study such as increasing joint level stiffness, changing reflex gains, and increasing cognitive focus. Additionally, the subtle differences between treadmill and overground walking may have influenced participant gait. For instance, treadmill walking keeps gait speed constant and removes the ability to arrest gait in the face of destabilization or significantly vary step length without coincident variation of cadence. When walking overground in everyday life, such options are available and provide a wider range of strategies to select from and greater maneuverability. Participants may have also altered their stepping behavior due to the gap between the treadmill belts and the limited width of the treadmill for forward progression. Finally, the small size of the perturbation and repeated exposure may have resulted in participants not considering it a major threat after the initial exposures.

### e. Conclusion

This study investigated the effects of underfoot perturbations during gait with varying levels of physical certainty, temporal certainty, and temporal affordance. Contrary to our hypotheses, greater certainty (temporal or physical) did not contribute to the adoption of more refined anticipatory kinematic gait strategies. Rather, participants waited to assess instantaneous body state during the perturbation and recovered stable gait by adjusting stepping behavior in the steps following, and greater physical certainty allowed individuals to recovery normal gait sooner. A priori knowledge related to physical certainty may promote greater success in paradigms aimed at improvement of locomotor stability.

## 5. Acknowledgments

The authors would like to thank Tyler Ho and Claire Rogers for their assistance with data collection and processing.

## 6. Competing Interests

No competing interests declared.

## 7. Funding

This work was supported by the Eunice Kennedy Shiver National Institute of Child Health and Human Development of the National Institutes of Health under award No. K12HD073945 (PCF). Opinions, interpretations, and conclusions are those of the author and are not necessarily endorsed by the funders.

## Abbreviations

CoM: Center of mass
vCoM: Velocity of the center of mass
xCoM: Extrapolated center of mass
CoP: Center of Pressure
BoS: Base of support
SW: Step width
MoS: Margin of stability
ML: Mediolateral
mTBI: Mild traumatic brain injury

## Notes

**Declarations of Interest:** None

### Competing Interest Statement

The authors have declared no competing interest.

