## Supplemental Data for "The effects of physical and temporal certainty on locomotion with discrete underfoot perturbations"

1.1 *Analysis of temporal and kinetic adjustments to stepping behavior before and after the perturbation*

Due to foot placement being the predominant balance control strategy during gait, our initial analyses focused solely on SW and MoS. However, healthy adults sometimes use alternative strategies during gait such as adjusting joint torque or the temporal sequence of stance and swing. To account for the possibility of individuals using such alternative strategies, we ran supplementary analyses examining stance time, swing time, and vertical impulse for steps both before and after the perturbation. These supplementary analyses include the same sample of participants as the primary analyses. Stance time was defined as the time from heel strike to toe off during each step while swing time was defined as the time from toe off to heel strike between each step. Vertical impulse was calculated by taking the integral of the vertical ground reaction force for each step.

1.2 *Interpretation of temporal and kinetic adjustments to stepping behavior*

Participants didn’t exhibit substantially altered stance times (Supplementary Figure 1) or swing times (Supplementary Figure 2) for steps before or after the perturbations were delivered. There was very little difference between conditions at each step, suggesting that neither affordance nor certainty had a large impact on the temporal dynamics of gait.

Vertical impulse did not appear to be influenced by condition; however, we did observe a consistent reduction in vertical impulse during the perturbation step relative to anticipatory and recovery steps (Supplementary Figure 3). A reduction in impulse could be caused by a reduction in stance time or a reduction in peak force. When considered with the stance time results above (Supplementary Figure 1), it appears that participants reduced the overall load on the perturbed limb to mitigate the magnitude of disruption caused by the perturbation.

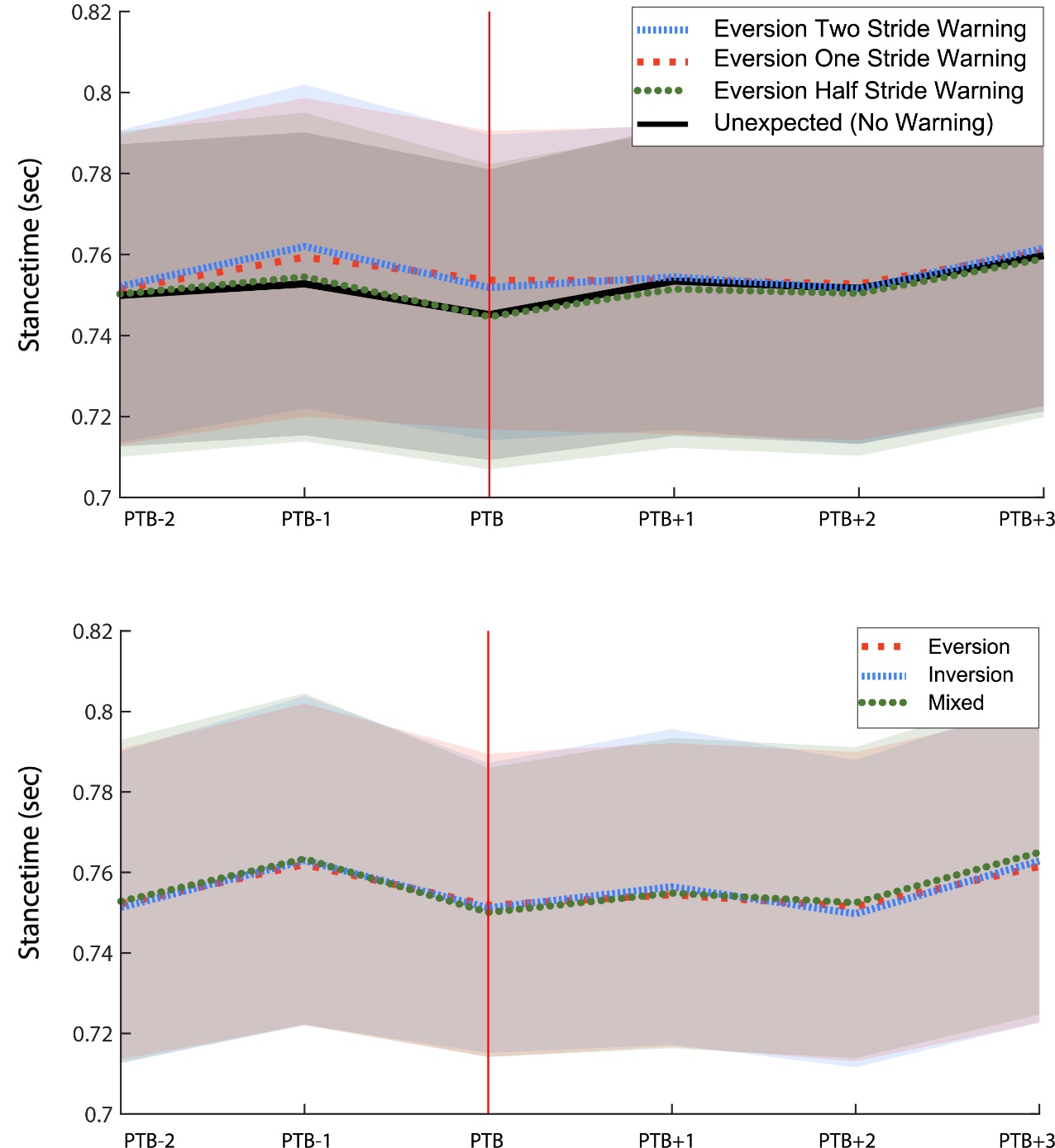

**Supplementary Figure 1:** Stance time for steps surrounding the perturbation. Conditions testing temporal affordance and temporal certainty are displayed in the top window and conditions testing physical certainty are displayed in the bottom window. Solid lines represent mean and shaded regions represent the 95% confidence intervals.

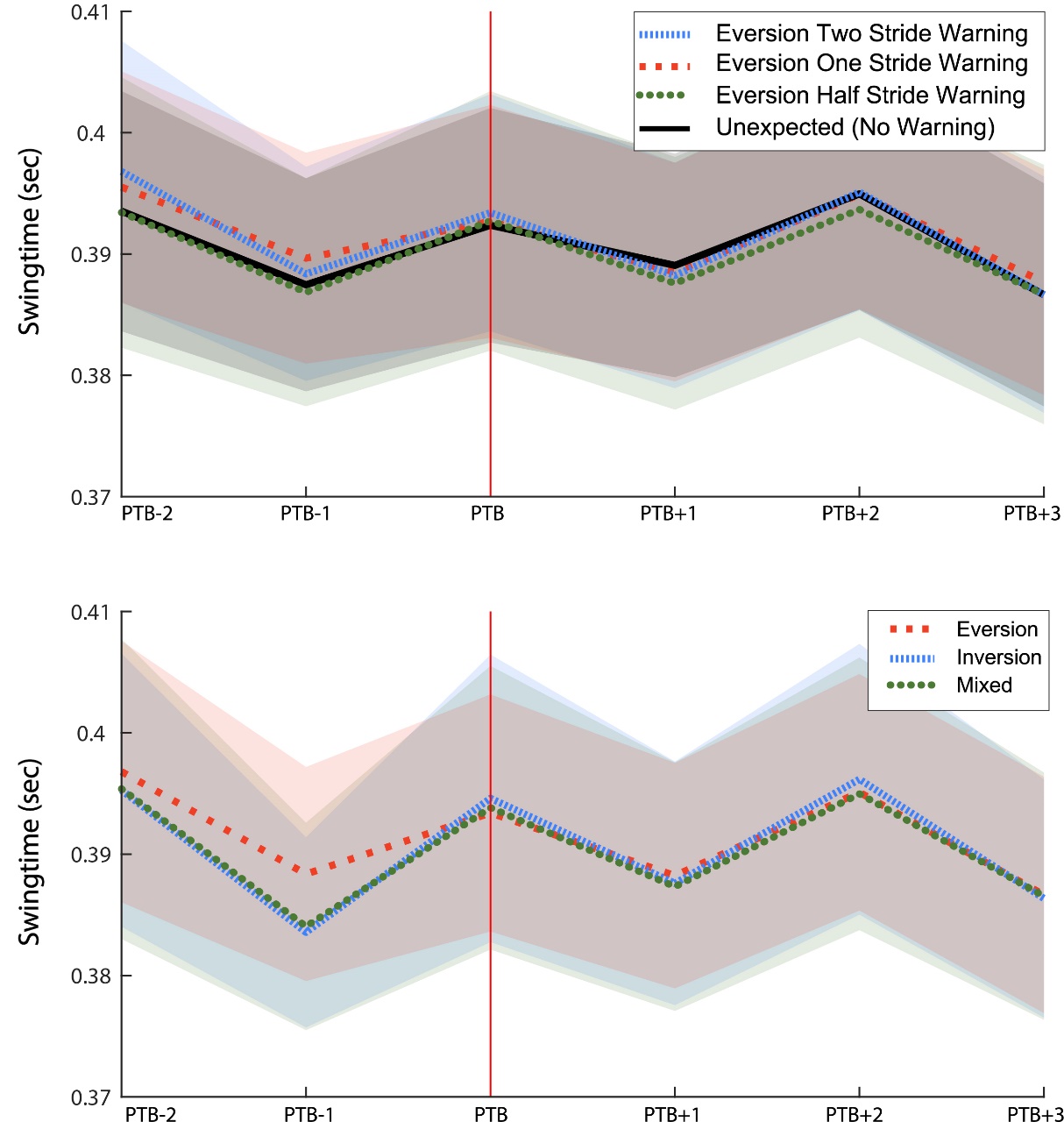

**Supplementary Figure 2:** Swing time for steps surrounding the perturbation. Conditions testing temporal affordance and temporal certainty are displayed in the top window and conditions testing physical certainty are displayed in the bottom window. Solid lines represent mean and shaded regions represent the 95% confidence intervals.

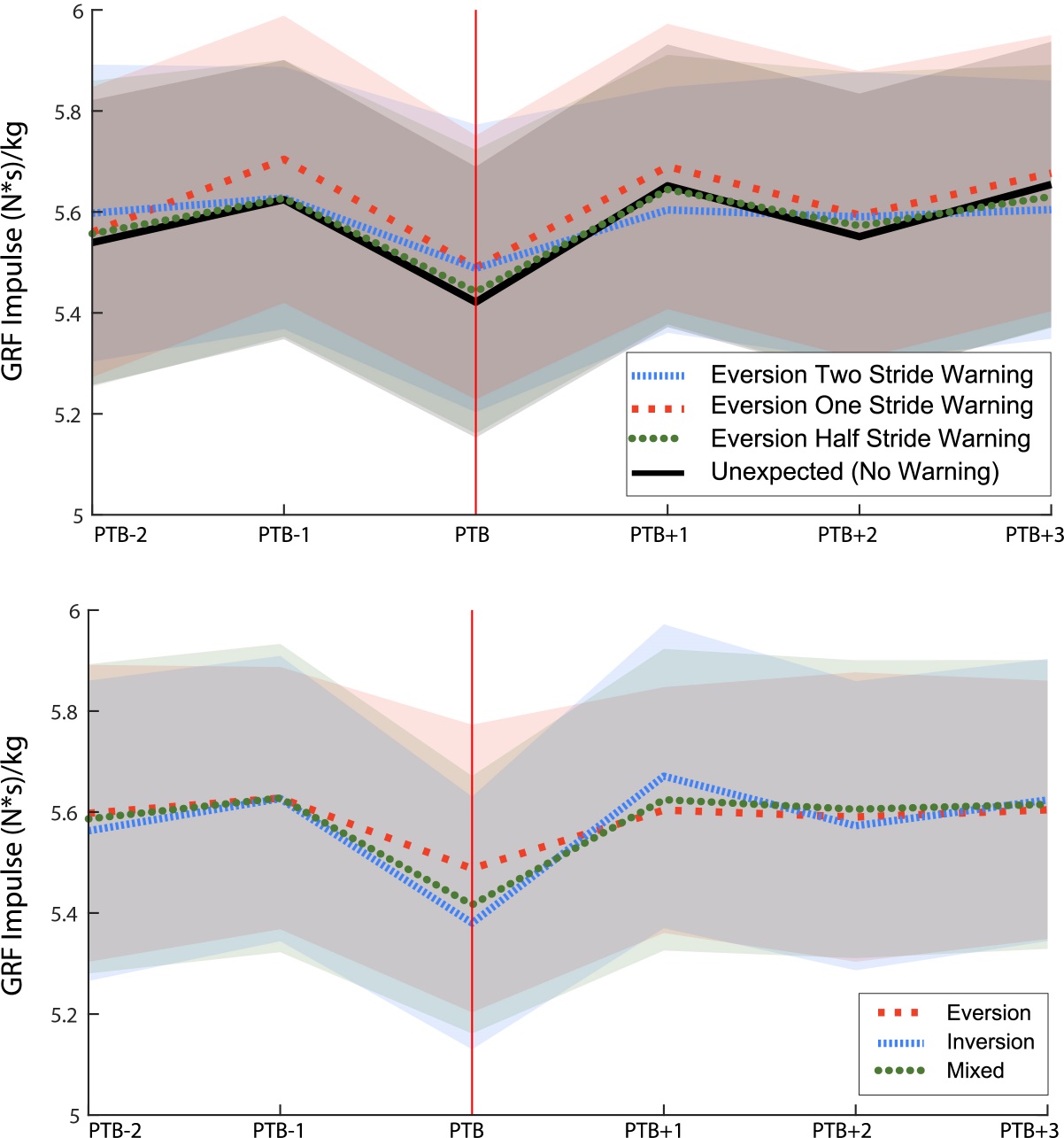

**Supplementary Figure 3:** Mass normalized vertical impulse for steps surrounding the perturbation. Conditions testing temporal affordance and temporal certainty are displayed in the top window and conditions testing physical certainty are displayed in the bottom window. Solid lines represent mean and shaded regions represent the 95% confidence intervals.
